# MinION sequencing of seafood in Singapore reveals creatively labelled flatfishes, confused roe, pig DNA in squid balls, and phantom crustaceans

**DOI:** 10.1101/826032

**Authors:** Jonathan K. I. Ho, Jayanthi Puniamoorthy, Amrita Srivathsan, Rudolf Meier

**Author notes:** First authors (Jonathan K. I. Ho, Jayanthi Puniamoorthy).

## Abstract

Food mislabelling is a growing world-wide problem that is increasingly addressed through the authentication of ingredients via techniques like mass spectrometry or DNA-sequencing. However, traditional DNA sequencing methods are slow, expensive, and require well-equipped laboratories. We here test whether these problems can be overcome through the use of Nanopore sequencing. We sequenced 92 single and 13 mixed-species samples bought in supermarkets and restaurants in Singapore which has a large and diverse seafood trade. We successfully obtained DNA barcodes for 94% and 100% of the single- and mixed-species products after correcting the numerous sequencing errors of MinION reads with a correction pipeline optimized for DNA barcodes. We find comparatively low levels of clear-cut mislabelling for single-species samples (7.6 %) while the rates are higher for mixed-species samples (38.5 %). These low rates are somewhat deceptive, however, because of the widespread use of vague common species names that do not allow for a precise assessment of the expected ingredients. With regard to the clearly mislabelled single-species products, higher-value products (e.g., prawn roe, wild-caught Atlantic salmon, halibut) are replaced with lower-value ingredients (e.g., fish roe, Pacific salmon, arrowtooth flounder) while more serious problems are observed for mixed-species samples. Cuttlefish and prawn balls repeatedly contained pig DNA and 100% of all mixed samples labelled as containing crustaceans (‘crab’, ‘prawn’, ‘lobster’) only yielded fish barcodes. We conclude that there is a need for more regular testing of seafood samples and suggest that due to speed and low-cost, MinION would be a good instrument for this purpose. We also emphasize the need for developing clearer labelling guidelines.

## 1. Introduction

In today’s globalised economy, seafood readily moves across borders. Fish caught in the Arctic and the Antarctic is served in restaurants on the equator, while scallops, oysters, and sea cucumbers harvested from the shores of North America satisfy the ever-increasing demand of consumers in East Asia. This increased demand has also led to the expansion of seafood farming worldwide. However, increased demand has also created incentives for seafood fraud via mislabelling. Such fraud is particularly common for fillets and heavily processed seafood products because they are not readily identifiable by eye (Boughattas, Le Fur, & Karoui, 2019; Carvalho, Palhares, Drummond, & Gadanho, 2017; Di Pinto et al., 2013; Giusti, Armani, & Sotelo, 2017; Veneza et al., 2018).

In recent years, seafood fraud and mislabelling have attracted much attention and the scope of the problem has become more apparent. This is partly because new technologies have made it easier to detect fraud. Most fraud appears driven by the desire to maximize profit because profit margins can be significantly increased by substituting expensive and desirable food species with less desirable and cheaper ones. For example, tilapia (*Oreochromis* spp.) or pangasius (*Pangasianodon hypophthalmus)* are occasionally sold as more expensive fish such as snapper or cod (Hu, Huang, Hanner, Levin, & Lu, 2018; Kappel & Schröder, 2015; Khaksar et al., 2015; Nagalakshmi, Annam, Venkateshwarlu, Pathakota, & Lakra, 2016; Pardo et al., 2018). Similarly, farmed Atlantic salmon (*Salmo salar*) is sold as wild-caught Pacific salmon (*Onchorhynchus* spp.) (Cline, 2012), and farmed rainbow trout (*Oncorhynchus mykiss*) as wild-caught brown trout (*Salmo trutta*) (Muñoz-Colmenero, Juanes, Dopico, Martinez, & Garcia-Vazquez, 2017).

But seafood mislabelling is sometimes more than “just” consumer fraud. It can also affect food safety when toxic or unpalatable species such as pufferfish or escolar enter the market by relabelling them as palatable species (Huang et al., 2014; Lowenstein, Amato, & Kolokotronis, 2009; Xiong et al., 2018). In addition, mislabelling frequently interferes with the conservation of species and populations when they are sold although they are protected by law (Almerón-Souza et al., 2018; Marko et al., 2004; Marko, Nance, & Guynn, 2011; Wainwright et al., 2018). Finally, an additional and underappreciated problem is that mislabelled food may contain ingredients that violate religious rules or ethical preferences, given that the consumption of some ingredients are disallowed or discouraged by specific religions.

The number of studies examining seafood fraud have increased greatly in recent years (Cawthorn, Baillie, & Mariani, 2018; Harris, Rosado, & Xavier, 2016; Pardo et al., 2018; Shehata, Bourque, Steinke, Chen, & Hanner, 2019). Several methods have been developed that are able to identify the ingredients of commercially sold seafood. This includes chromatographic, spectroscopic, proteomic and genetic methods. Protein-based methods are particularly well-established for the identification of commonly traded fish species. They were the first molecular method for identifying ingredients of seafood products to species and they remain very popular in the form of mass spectroscopy (MS) which has the advantages of being fast and comparatively low-cost (Black et al., 2017; Mazzeo & Siciliano, 2016; Stahl & Schröder, 2017; Wulff, Nielsen, Deelder, Jessen, & Palmblad, 2013). However, identification requires comprehensive databases of MS profiles for the traded seafood, which are difficult to develop for rare species, heavily processed samples, and samples consisting of mixtures of multiple species.

For these reasons, genetic methods have recently received more attention. They have high accuracy and specificity (Haynes, Jimenez, Pardo, & Helyar, 2019) and benefit from the large number of seafood species that have been characterized with DNA barcodes. Genetic testing of seafood ingredients generally relies on the standard DNA barcode for animals; i.e., an approximately 650bp long segment of the mitochondrial cytochrome oxidase I (COI) gene. Reference sequences for this barcode are available for a large number of commercially traded species. This has the advantage that most sequences obtained from seafood products can be assigned to species or species-groups. In addition, mixed- and heavily processed samples can still be characterized because they still contain trace amounts of DNA. However, DNA barcodes are only slowly becoming popular for food authentication because of the comparatively high cost of sequencing when they are obtained with Sanger sequencing (e.g., cost per barcode at the Canadian Centre for DNA barcoding is USD 17: http://ccdb.ca/pricing/). Furthermore, Sanger sequencing does not allow for sequencing products that contain signals from multiple species. Fortunately, these problems can be overcome by using new sequencing methods that are often collectively referred to as Next-Generation sequencing (NGS) or High Throughput Sequencing technologies (HTS). DNA barcodes obtained on platforms such as Illumina, Ion Torrent, and PacBio have been used for food authentication (Carvalho et al., 2017; Giusti et al., 2017; Xing et al., 2019), but they have several disadvantages. The equipment and maintenance cost for Illumina and PacBio instruments are so high that these sequencers are mostly found in sequencing centres that have fairly long turnaround times for submitted samples. In addition, due to the high cost of flowcells, the cost per DNA barcode is high unless thousands of products are sequenced at a time (Ho, Foo, Yeo, & Meier, 2019; Kutty et al., 2018; Srivathsan et al., 2018; Wang, Srivathsan, Foo, Yamane, & Meier, 2018; Yeo, Puniamoorthy, Ngiam, & Meier, 2018).

Fortunately, these issues can now be addressed with Oxford Nanopore sequencing which is implemented on small and portable MinION™ sequencers. This technology could potentially have three key advantages for food authentication. Firstly, the sequencer and the flowcells are sufficiently inexpensive to make them suitable for routine testing in many laboratories and regulatory agencies. In addition, the cost per sample is quickly dropping because recent advances in bioinformatic pipeline now allow for obtaining up to 3500 barcodes on a single standard flowcell (Srivathsan et al., 2018). Furthermore, even less expensive flowcells with lower capacity have become available that will be suitable for processing a few hundred samples. Secondly, obtaining barcodes with MinION requires minimal lab equipment and the data can even be obtained under difficult field conditions ranging from hot, humid tropical rainforest (Pomerantz et al., 2018; Schilthuizen et al., 2019) to freezing Antarctic habitats (Johnson, Zaikova, Goerlitz, Bai, & Tighe, 2017). This is why MinION is not only suitable for rapid species discovery (Schilthuizen et al., 2019; Srivathsan et al., 2018, 2019) but also for identifying species under challenging circumstances (Blanco et al., 2019; Parker, Helmstetter, Devey, Wilkinson, & Papadopulos, 2017; Pomerantz et al., 2018). Lastly, MinION devices generate data within minutes of loading a flowcell and allow for data collection in real-time. Given all these advantages, one may ask why MinION sequencers are not the default for food authentication with DNA sequences. Presumably, the main reason is the high sequencing error rate of 10-15% (Wick, Judd, & Holt, 2019), but fortunately these errors can now be effectively corrected using a range of new bioinformatics pipelines that are optimized for obtaining animal barcodes with nanopore sequencers (Maestri et al., 2019; Srivathsan et al., 2018).

Currently, MinION sequencing has apparently only been used in one study addressing seafood authentication (Voorhuijzen-Harink et al., 2019). It compared the accuracy of MinION results with those of other high-throughput sequencing techniques and found them to be similar. However, the study did not examine seafood products sold commercially and only examined two artificially mixed samples. The study also predated recently improved bioinformatics pipelines for obtaining DNA barcodes with MinION (Srivathsan et al., 2019). These limitations are here overcome by studying >100 samples of seafood sold in Singapore. The data are analysed using these newly developed techniques and we analyse both single- and mixed-species samples using two different primer pairs. Our study furthermore contributes to the still very limited amount of information available on the prevalence of seafood fraud in Southeast Asia (Labrador, Agmata, Palermo, Follante, & Pante, 2019; Sarmiento, Pereda, Ventolero, & Santos, 2018; Sultana et al., 2018; Too, Adibah, Danial Hariz, & Siti Azizah, 2016; Tran, Nguyen, Nguyen, & Guiguen, 2018). Note that Singapore is a good area for developing seafood authentication methods because it is a very large seafood market. The city state imported 129,439 tonnes of seafood in 2017 (70% being fish and 30% being other seafood), while only producing 6,498 tonnes (91% fish and 9% other seafood)(Agri-Food and Veterinary Authority, 2018). Average per capita consumption is an estimated 21 kg (71% fish and 29% other seafood), which is slightly above the world average of 20.5 kg (FAO, 2018). Overall, Singapore residents obtain nearly 30% of their animal protein from seafood, yet seafood products purchased in Singapore have only been included in two authentication studies. The first established the identity of commercially sold ‘snappers’ in six English-speaking countries (Cawthorn et al., 2018) while the second examined the species identity of commercially available elasmobranchs. The latter revealed that in Singapore, shark meat (Carcharhinidae) was being sold as Indian threadfin (*Leptomelanosoma indicum*) (Wainwright et al., 2018).

## 2. Materials and Methods

### 2.1 Sample collection

We obtained 105 samples of fresh and frozen seafood from 6 supermarkets (Table 2, Table 3, Table 5) and 2 seafood restaurants (Table 4) in Singapore. All samples were purchased in the first week of May 2018, and each location was visited only once. The products were divided into two categories, single-species products (e.g., frozen fillets) and mixed-species products (e.g., fish or squid balls). All samples did not undergo any cooking or processing after purchase. We tested 92 single-species products (21 from restaurants and 71 from supermarkets) and 13 mixed-species products (all from supermarkets).

**Table 1:** DNA barcoding data obtained with MinION

|  | COI full length barcode | COI minibarcode |
| --- | --- | --- |
| Number of reads demultiplexed | 158,329 | 91,901 |
| Coverage (Average/median/range) | 1508/773/13-14,211 | 875/403/12-6071 |
| Number of single-species sample barcodes | 70 | 68 |
| Number of mixed-species samples with barcodes | 14 | 15 |

**Table 2.** Mislabelled single-species samples of seafood products obtained from supermarkets in Singapore. Ambiguities in identification are separated by slashes.

| Seafood sold/marketed as: (expected species) |  | ID of barcode (best match %) |
| --- | --- | --- |
| FM027 | Premium Halibut fillet (*Hippoglossus* sp.) | **313 bp:** *Atherethes stomias* (100%)<br>**658 bp:** *Atherethes stomias* (100%) |
| FM041 | Wild caught Atlantic Salmon boneless fillet (*Salmo salar*) | **313 bp:** *Oncorhynchus keta* (100%)<br>**658 bp:** Unsuccessful |
| FM049 | Sole fillet (*Soleidae* sp.) | **313 bp:** *Psettodes erumei* (100%)<br>**658 bp:** Unsuccessful |
| FM072 | Premium Halibut fillet (*Hippoglossus* sp.) | **313 bp:** *Atherethes stomias* (100%)<br>**658 bp:** *Atherethes stomias* / *A. evermanni* (100%) |
| FM073 | Wild-caught Atlantic Salmon boneless fillet (*Salmo salar*) | **313/658 bp:** *Oncorhynchus keta* (100%) |
| FM074 | Crab leg (*Brachyura* sp.)* | **313 bp:**<br>*Nemipterus mesoprion* / *N. randalli* (100%)<br>**658 bp:**<br>*Epinephelus diacanthus* (99.85%); *Nemipterus mesoprion* / *N. randalli* (100%);<br>*Panna microdon* (99.85%); *Priacanthud hamrur* / *P. prolixus* (100%); *Scolopsis taenioptera* (100%) |
| FM077 | Prawn Roe (Penaeidae sp.) | **313/658 bp:** *Mallotus villosus* (100%) |

\* This product was expected to be single-species but analysis revealed multiple species (separated by semicolons)

**Table 3:**
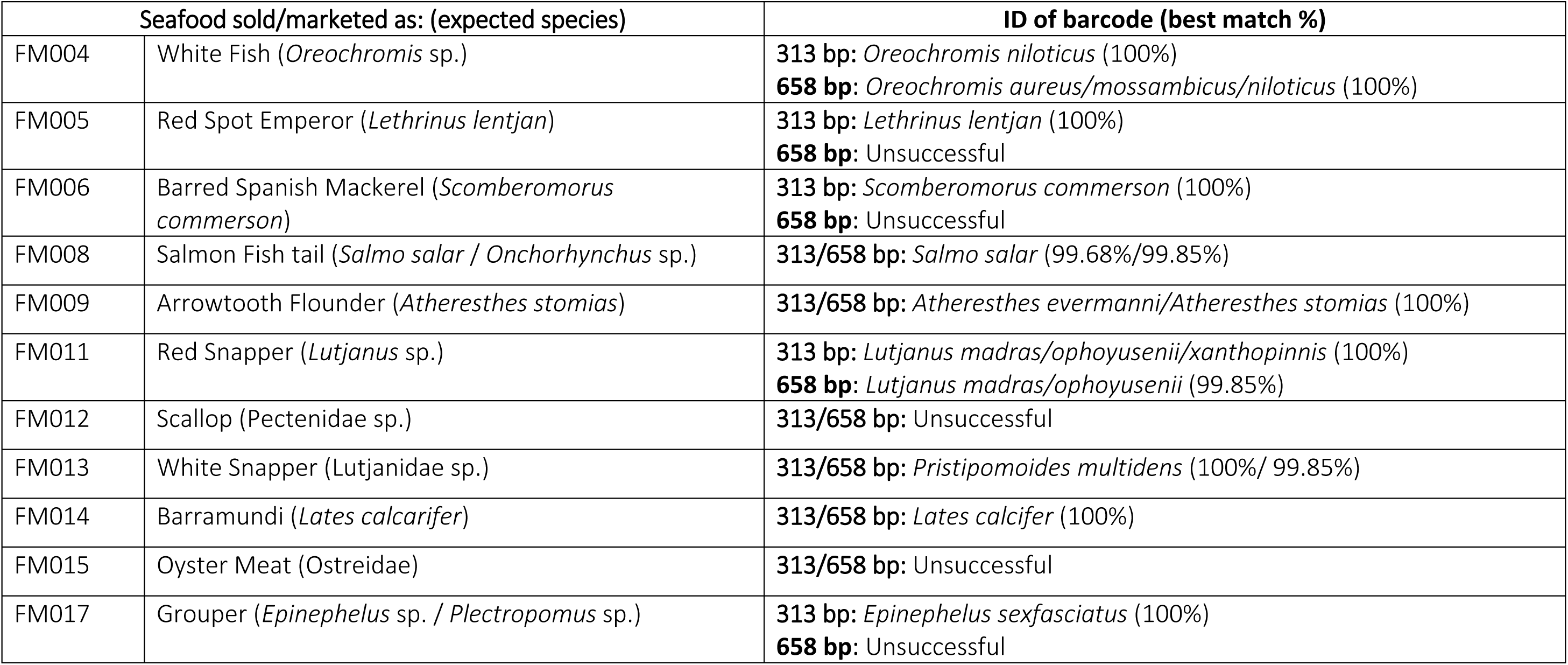
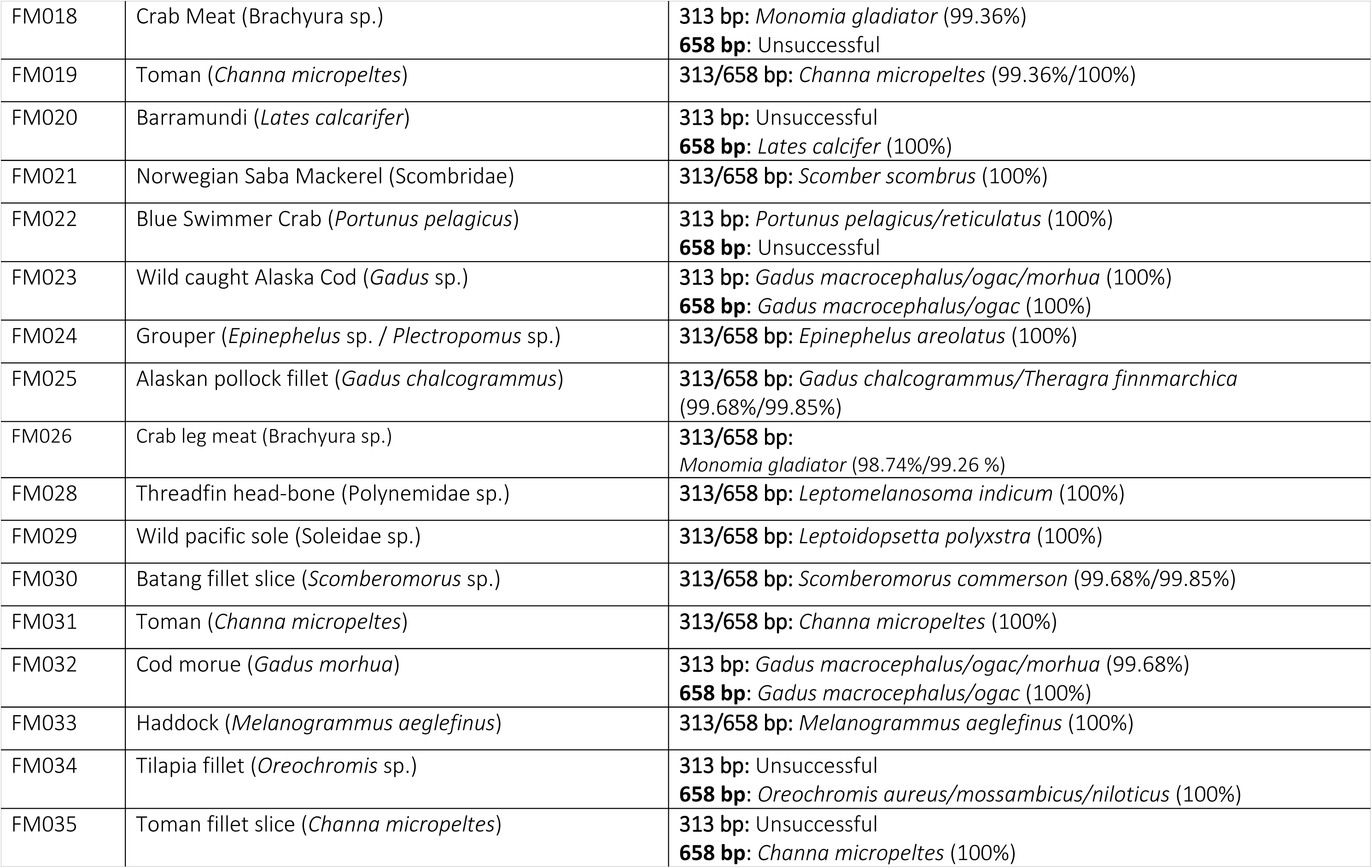
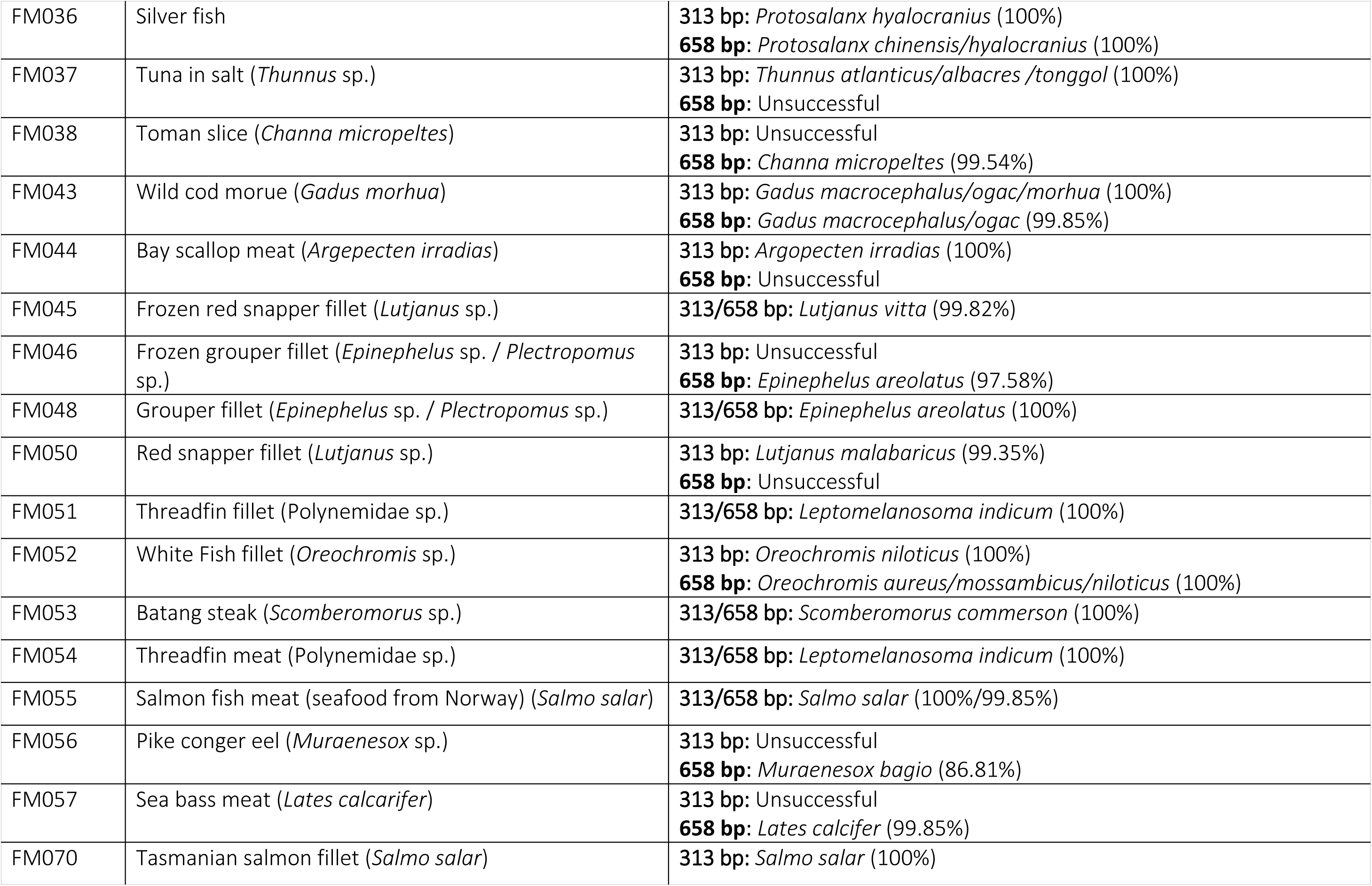
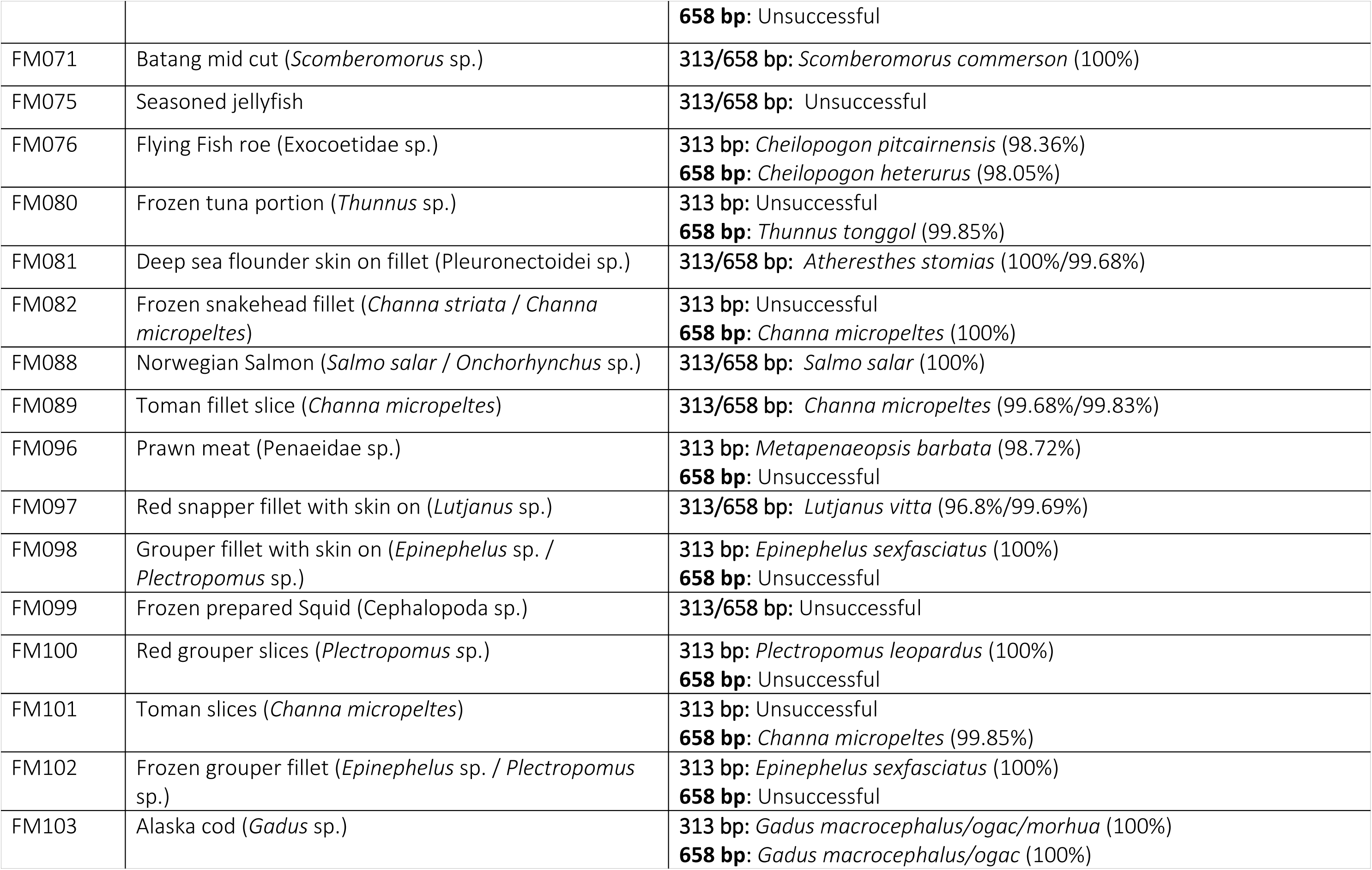
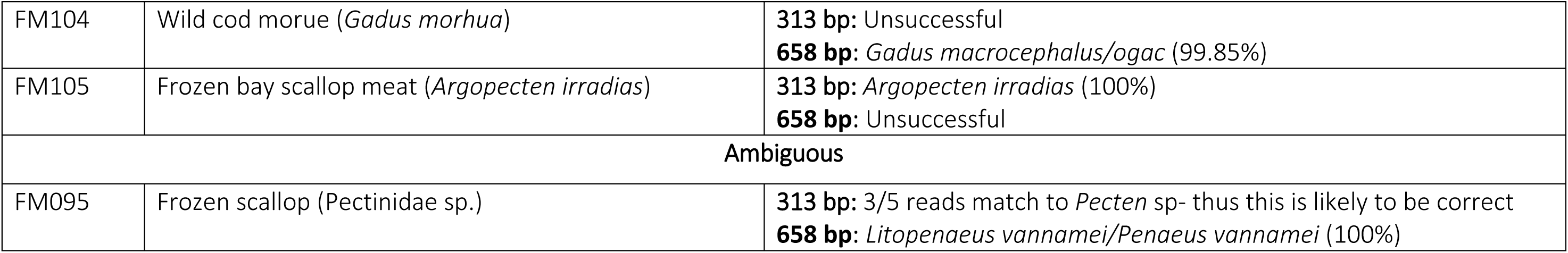
Correctly labelled single-species samples of seafood products obtained from supermarkets in Singapore. Ambiguities in identification are separated by slashes.

**Table 4:** Single-species samples of seafood products obtained from restaurants in Singapore. Ambiguities in identification are separated by slashes.

| Seafood sold/marketed as: (expected species) |  | ID of barcode (best match %) |
| --- | --- | --- |
| FM058 | Eel Sushi Roll (*Anguilla* sp.) | **313 bp:** Unsuccessful<br>**658 bp:** *Anguilla anguilla* (100%) |
| FM059 | Salmon Roll (*Salmo salar* / *Onchorhynchus* sp.) | **313/658 bp:** *Onchorhynchus keta* (100%/99.84%) |
| FM060 | Scallop (*Pectenidae* sp.) | **313/658 bp:** Unsuccessful |
| FM061 | Red Sea Bream/Amberjack (*Pagrus* sp. / *Pagellus* sp. / *Seriola* sp.) | **313 bp:** *Pagrus major/auratus* (100%)<br>**658 bp:** Unsuccessful |
| FM062 | Yellow Fin Tuna (*Thunnus albacares*) | **313 bp:** Unsuccessful<br>**658 bp:** *Thunnus albacares* (99.85%) |
| FM063 | Red Sea Bream/Amberjack (*Pagrus* sp. / *Pagellus* sp. / *Seriola* sp.) | **313/658 bp:** *Seriola dumerili* (100%/99.85%) |
| FM064 | Northern Spot Prawn (*Pandalus platyceros*) | **313 bp:** Unsuccessful<br>**658 bp:** *Pandalus borealis* (100%) |
| FM065 | Salmon (*Salmo salar* / *Onchorhynchus* sp.) | **313/658 bp:** *Salmo salar* (100%/99.69%) |
| FM066 | Eel Sushi (*Anguilla* sp.) | **313/658 bp:** *Anguilla japonica/anguilla* (100%/99.85%) |
| FM067 | Codroe with Kelp (*Gadus morhua*) | **313 bp:** *Gadus macrocephalus/ogac/morhua* (100%) |
|  |  | **658 bp:** *Gadus macrocephalus/ogac* (99.85%) |
| FM068 | Yellowfin tuna (*Thunnus albacares*) | **313 bp:** *Thunnus atlanticus/albacres/tonggol* (100%)<br>**658 bp:** *Thunnus albacares* (100%) |
| FM069 | Golden Cuttlefish (*Sepia esculenta/ Sepia elliptica*) | **313 bp:** *Sepia prashadi/pharaonis/ramani* (99.68%)<br>**658 bp:** *Sepia pharaonis* (99.84%) |
| FM084 | Salmon (*Salmo salar / Onchorhynchus* sp.) | **313/658 bp:** *Salmo salar* (100%/99.69%) |
| FM085 | Swordfish (*Xiphias gladius*) | **313/658 bp:** *Xiphias gladius* (100%) |
| FM086 | Bluefin tuna/Skipjack Tuna (*Thunnus* sp. / *Katsuwonus pelamis*) | **313 bp:** *Thunnus obesus/atlanticus/albacares* (100%)<br>**658 bp:** *Thunnus obesus* (99.85%) |
| FM087 | Bluefin tuna/Skipjack Tuna (*Thunnus* sp. / *Katsuwonus pelamis*) | **313/658 bp:** *Katsuwonus pelamis* (100%/99.85%) |
| FM090 | Tara kirmi (cod fish fillet) (*Gadus* sp.) | **313/658 bp:** *Gadus macrocephalus/ogac* (100%) |
| FM091 | Kajiki kirmi (swordfish fillet) (*Xiphias gladius*) | **313/658 bp:** *Xiphias gladius* (100%) |
| FM092 | Gindara kirmi (black cod fillet) (*Anoplopoma fimbria*) | **313/658 bp:** *Anoplopoma fimbria* (100%) |
| FM093 | Buri kirmi (Yellowtail fillet) (*Seriola quinqueradiata*) | **313 bp:** Unsuccessful<br>**658 bp:** *Seriola lalandi/quinqueradiata* (100% - 99.66%) |
| FM094 | Seasoned cod roe (*Gadus morhua*) | **313/658 bp:** *Gadus chalcogrammus/Theragra finnmarchica* (100%/100% - 99.23%) |

**Table 5:**
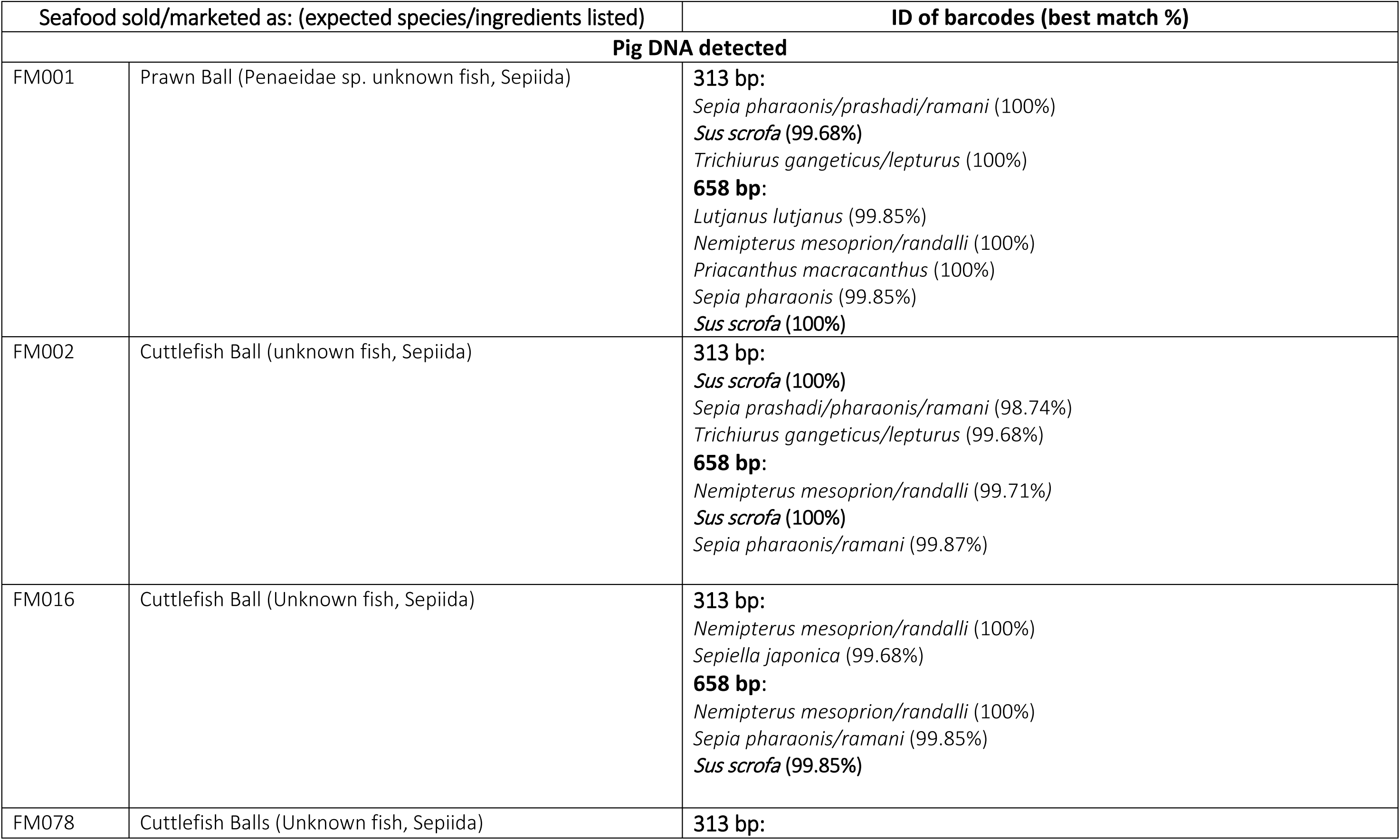
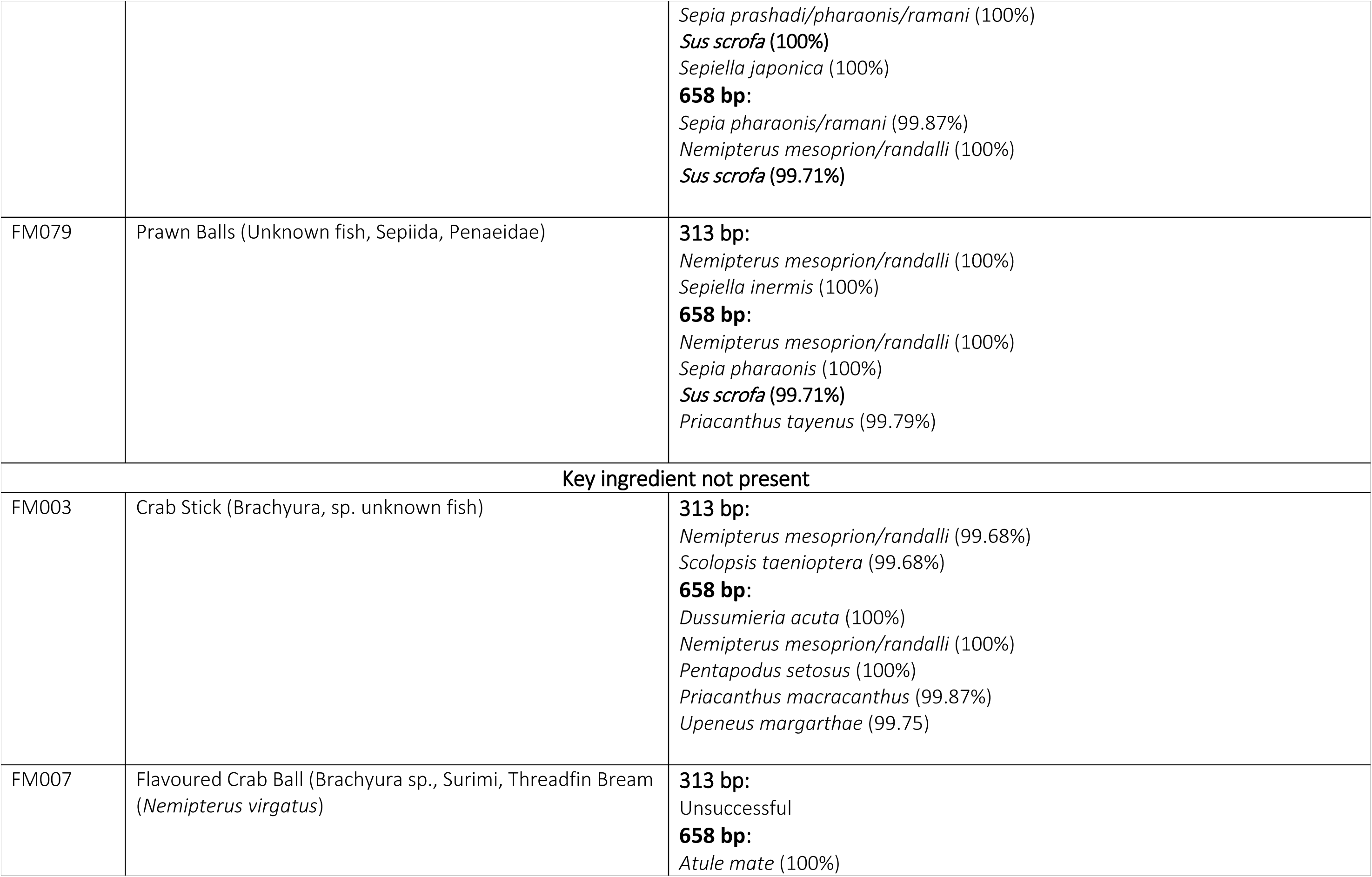
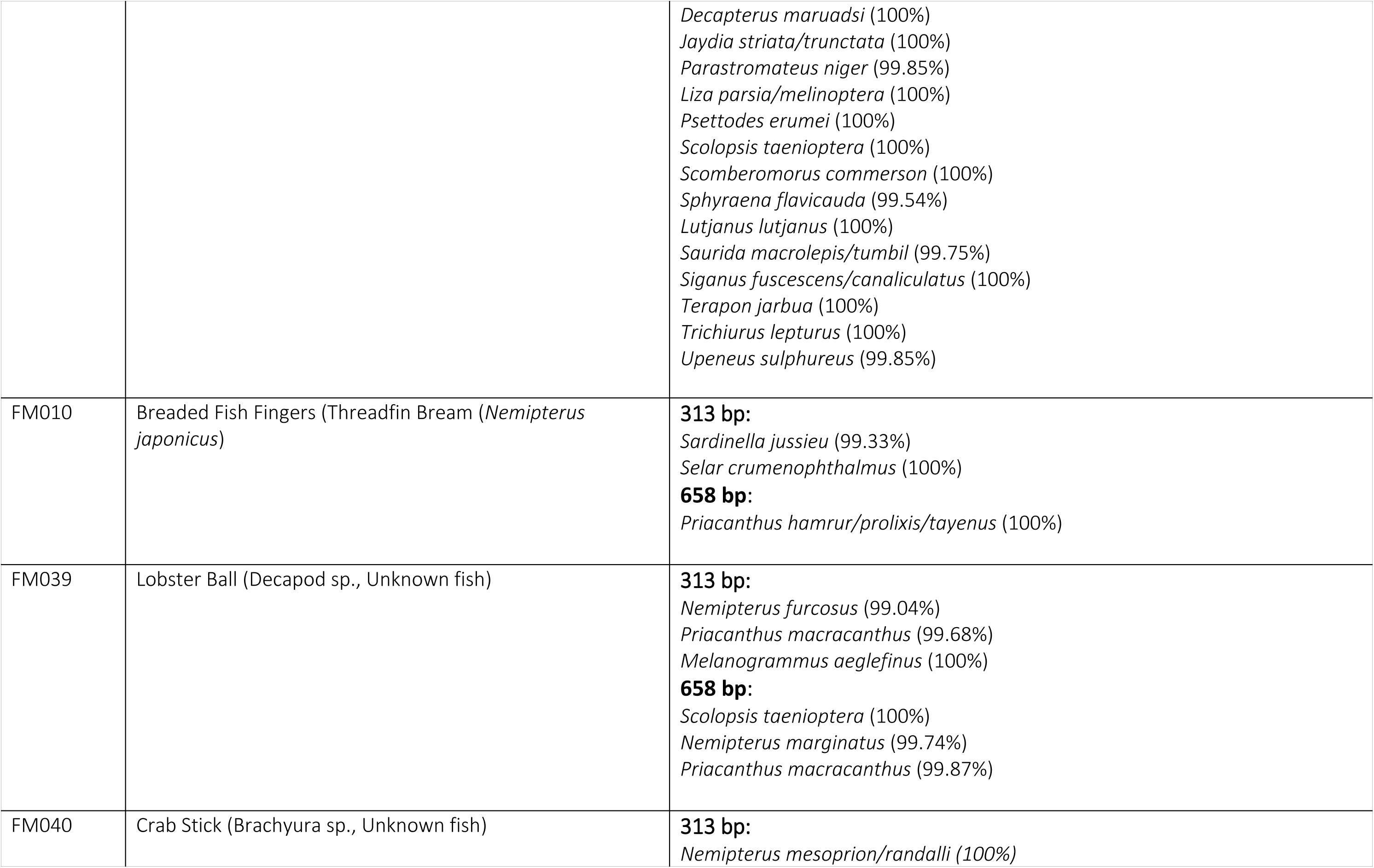
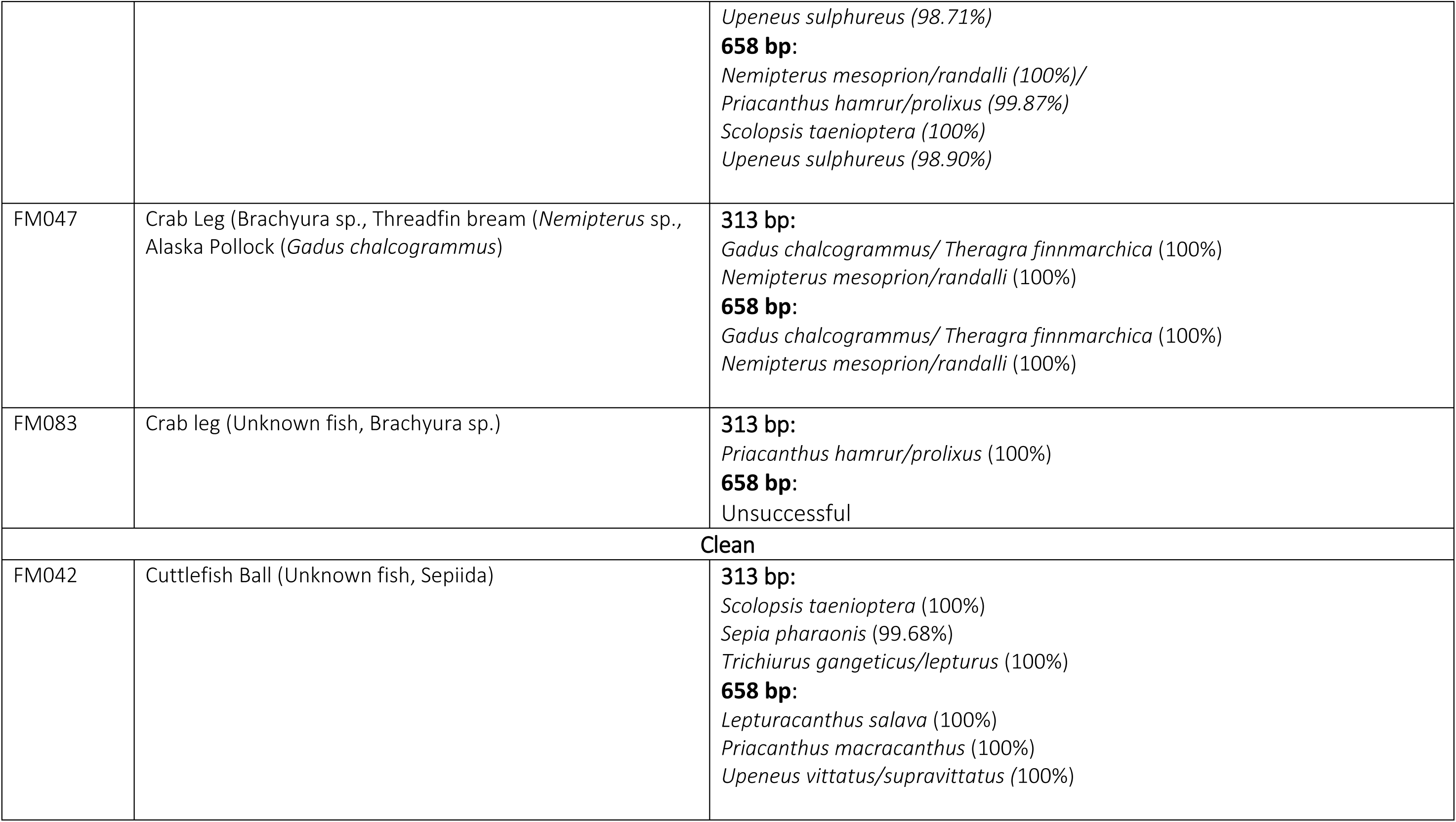
Mixed-species samples obtained from supermarkets in Singapore. Ambiguities in identification are separated by slashes.

### 2.2 DNA extraction and PCR

DNA extraction was conducted using an automated extraction system (Bioer Automatic Nucleic Acid Purification system) using MagaBio plus Tissue Genomic DNA purification kit using the manufacturer’s protocols. Afterwards, we amplified two barcodes that differed in length. In order to obtain full length DNA barcodes, we used a COI-3 primer cocktail (C_FishF1t1–C_FishR1t1, (Ivanova, Zemlak, Hanner, & Hebert, 2007)), while a shorter mini-barcode (313 bp) was obtained using m1COlintF: 5’-GGWACWGGWTGAACWGTWTAYCCYCC-3’ (Leray et al., 2013) and a modified jgHCO2198: 5’-TANACYTCNGGRTGNCCRAARAAYCA-3’ (Geller, Meyer, Parker, & Hawk, 2013). In order to multiplex a large number of samples in a single MinION run, we adopted a tagged amplicon strategy (Meier et al. 2016) where each primer was tagged with a 13-bp unique sequence at the 5’ end of the primer. Eleven forward and ten reverse tagged primers allowed for the amplification of 110 products using a dual-indexing strategy. For this study we used the tags developed by Srivathsan et al. (2019, F: HL001-HL011, R: HL001-HL010) and the PCR conditions for all amplifications were as follows: 8 μl Mastermix (CWBio), 7.84 μl molecular grade H_2_O, 0.16 μl of 25mM MgCl_2_, 1 μl of 1 mg/ml BSA, 1 μl of each primer, and 1 μl of sample DNA. The PCR conditions were 5 min initial denaturation at 94°C followed by 35 cycles of denaturation at 94°C (1 min), 47°C (2 min), 72°C (5 min), followed by final extension of 72°C (5 min). PCR products were pooled in equal volumes for library preparation and MinION sequencing. The libraries were prepared using SQK-LSK108 kit as per instructions using 1 µg of starting DNA. The only modification to the protocol recommendation by the manufacturer was the use of 1X Ampure beads for clean-up instead of the customarily recommended 0.4X. Sequencing was carried out using MinION R9.4.1 over 24 hours.

### 2.3 Bioinformatics

The nanopore reads were base-called in real-time using MinKNOW. The resulting fastq file was converted to a fasta file and the data were processed using *miniBarcoder* (Srivathsan et al., 2018, 2019). In short, the reads were split into two sets based on lengths (1) 300-600 bp and (2) >600 bp. The first read set was demultiplexed to obtain sequences corresponding to the COI minibarcode while the second read set included the reads pertaining to the full-length barcode. For this set, we first demultiplexed the reads using one pair of primers (FishF2_t1 and FishR2_t1) that were then removed from the read set. Next we used the second pair of primers (VF2_t1-FR1d_t1) for demultiplexing the remaining reads in the second set. The average coverage for two combinations was >1000 X (median 770X) with all specimens having >10X coverage. Hence, we did not proceed to recover additional reads by demultiplexing the remaining primer combinations.

A bioinformatics pipeline for single-species barcodes from sets of reads developed by Srivathsan et al. (2018, 2019) was used here. Briefly, it first obtains a “MAFFT barcode” by aligning the reads using MAFFT (Katoh & Standley, 2013) and obtaining a majority rule consensus with subsequent removal of gaps. These MAFFT barcodes are further corrected using RACON (Vaser, Sovic, Nagarajan, & Sikic, 2017) to generate a second set of consensus barcodes. The MAFFT and RACON barcodes are then corrected for indel errors based on amino-acid translations. Lastly these barcode sets are consolidated to obtain final barcodes. For mixed species products, we modified the bioinformatics procedures. For each sample, the demultiplexed reads were matched by BLAST to GenBank (e-value threshold of 1e-5). The BLAST matches were then parsed using *readsidentifier* (Srivathsan, Sha, Vogler, & Meier, 2015) to summarize the taxonomy using the Lowest Common Ancestor approach and retaining only the best scoring matches. Read sets were grouped by genus, and the abovementioned pipeline was used to obtain a consensus barcode for each genus specific read set. This approach was also applied to read sets for samples for which we failed to get clean barcodes using the single-species approach. This is because bacterial signals can be co-amplified with a seafood product, and a clean barcode sequence can only be obtained after the removal of the bacterial reads.

All barcode sequences were matched by BLAST to NCBI NT database and the 50 best matches were retrieved. These were aligned with the barcode datasets using MAFFT and queried with SpeciesIdentifier (Meier, Shiyang, Vaidya, Ng, & Hedin, 2006) to find the best matching sequence.

## 3. Results and Discussion

### 3.1. Amplification success

The use of two different sets of primers amplifying the full-length and a mini-barcode of 313bp length allowed us to obtain sequences for 87/92 (94.5%) of the single-species and 13/13 (100%) of the mixed-species products. These barcodes were derived from 158,329 short and 91,901 long nanopore reads that were successfully demultiplexed into read sets representing the different amplicons. This overall high success rate is due to combining the data for both amplicons. We obtained mini-barcodes for 72 and full-length barcodes for 70of the 92 single-species samples, but only 55 samples (60%) have data for both. We thus strongly recommend the use of different primer sets in order to increase the overall success rates. The usage of two different PCR reactions furthermore helps with overcoming potential primer biases and allows for cross-validation. For example, one sample (FM095) was expected to contain frozen scallop but a prawn DNA barcode was obtained when using the full-length primer cocktail. In contrast, the mini-barcode reads revealed the expected scallop signal. Once this sample is excluded from the analysis, our total success rate for single-species products is 93.4%, since no other samples failed this cross validation. Note that for mixed products, the success rates were higher than for single-species products. This applies to both sets of primers (12 of 13 samples had at least one sequence successfully barcoded) and was surprising because we had expected that such samples would be more difficult to sequence. By matching the barcodes to publicly available reference sequences, we classified seven single-species samples (7.6%: Table 2) and five mixed species samples (Table 5) as being clearly mislabelled. However, we submit that an additional seven mixed-species samples are borderline mislabelled and the labelling could be considered fraudulent if stricter rules were applied to the equivalence of scientific and common names.

### 3.2. Identification of seafood samples

Several of the clear-cut cases of mislabelling involved flatfish for which about 40% of all single-species samples were affected (3 out of 7). This includes two cases of halibut (*Hippoglossus* sp.) being substituted by arrowtooth flounder (*Atheresthes stomias*) and one sample of sole (*Solea* sp.) being substituted by Indian halibut (*Psettodes erumei*). Similar to cases reported elsewhere in the literature, salmon were also targeted with two samples of chum salmon (*Onchorhynchus keta*) being sold as wild-caught Atlantic salmon (*Salmo salar*). We also found that one sample of capelin roe (*Mallotus villosus*) was sold as prawn roe. Arguably, the most serious case of mislabelling for a multi-religious society like Singapore involved pig DNA in cuttlefish and prawn balls. We initially suspected lab contamination, but the same seafood brand repeatedly yielded pig DNA in five samples which were bought at different times and places. Pig DNA was also consistently amplified by both primer sets and were not found in any of the other seafood samples. This ingredient in a seafood product is a serious problem given that many consumers avoid pork for religious, ethical, or health reasons (e.g., allergies). Fortunately, the samples were not labelled as *halal* or *kosher*, but such cases do highlight the need for regular testing of heavily processed, multi-species seafood samples. Note that a similar case of pig DNA in seafood balls had also recently been reported from the Philippines (marketed as fish, squid, or shrimp balls). These seafood balls also included chicken meat (Sarmiento et al., 2018).

In most mislabelling cases, the substituted product was less valuable than the species indicated on the label. For example, halibut is a more highly valued fish compared to arrowtooth flounder, which tends to develop a soft and mushy texture when cooked (Greene & Babbitt, 1990). Arrowtooth flounder is found throughout the Eastern Pacific, from the Bering Sea to the coast of Baja California. Historically, it was not targeted by commercial fisheries because it was considered unpalatable, but new technology and population declines of other species have led to the exploitation of arrowtooth flounder populations (Grandin & Forrest, 2017). However, this does not change the fact that arrowtooth flounder can at best be considered a ‘low-value’ or even ‘nuisance’ species (Kasperski, 2016). Yet, it is starting to regularly show up in mislabelling studies, with recent cases reported from Brazil (Carvalho, Palhares, Drummond, & Frigo, 2015) and China (Xiong et al., 2016). We submit that explicit regulation is needed that requires that arrowtooth flounder be labelled as such. In addition, fast detection techniques targeting this species should be developed.

Not surprisingly, other cases of mislabelling involved salmon. “Wild-caught” Atlantic salmon (*Salmo salar*) was found to be chum salmon (*Onchorhynchus keta*). The latter species usually commands a lower price than wild-caught king or coho salmon (*O. tshawytscha*; *O. kisutch*) (Alaska Department of Fish and Game, 2018). This is presumably due to the fact that the commercial fishery for wild Atlantic salmon has now virtually collapsed due to significant population declines. Worldwide, the mislabelling of salmon usually involves farmed *S. salar* labelled as wild caught *Onchorhynchus* sp. or less valuable species of *Onchorhynchus* being substituted by more valuable ones (Cline, 2012; Muñoz-Colmenero et al., 2017; Warner et al., 2015). It appears that Singapore’s case of *O. keta* being labelled as “wild-caught” *S. salar* is a new addition to the numerous mislabelling problems in *Salmo* and *Onchorhynchus*.

Many mixed-species products were labelled as ‘crab’, ‘prawn’, or ‘lobster’ sticks or balls. Only fish were listed as ingredients in 6 out of 8 mixed-species samples while two more explicitly listed shrimp meat or prawn powder in addition to fish in their ingredients. However, we were unable to find any crustacean DNA in all eight samples. Fish DNA was abundant and we suspect that overall, many of these products do not include any or only minuscule amounts of crustacean tissues. One additional sample, which was simply labelled ‘crab legs’ without any ingredient list and was treated as a single-species product, proved to only contain fish DNA as well. One way or another, we submit that the average consumer would consider extremely low proportions of crustacean protein to be unacceptable given that the label highlights the crustacean component (‘crab’, ‘prawn’, ‘lobster’). This is in contrast to cuttlefish balls which usually contained cephalopods, usually from the cuttlefish genus *Sepia*. We suggest that this ‘creative labelling’ misleads consumers because the main product label indicates crustacean content and the fine print needs to be examined in order to determine that the product does not actually contain crustaceans. Note that the lack of crustacean signal is not due to primer biases because we used a mini-barcode primer mix that that is known to amplify a wide range of marine invertebrates; i.e., we would have expected to find crustacean DNA if it had been there.

### 3.3. Implications and suggestions

Overall, our study suggests that the level of clear-cut mislabelling of seafood products in Singapore is not particularly high when compared to results from other Southeast Asian countries. Studies from Malaysia, Vietnam and the Philippines found levels of mislabelling to be around 60% (Sarmiento et al., 2018; Sultana et al., 2018; Tran et al., 2018) with the only outlier study being by Too et al., (2016) who only detected seafood fraud in 16% of the tested seafood products in Malaysia. Unfortunately, establishing a baseline for overall levels of seafood mislabelling in the region is difficult because the studies are not directly comparable due to differences in methodology and sampling criteria. Hence, the next step for understanding and reducing the problem would be developing standardised sampling and analysis criteria. Sampling criteria could be the sales volume of a product (e.g., high-demand species like salmon, grouper, or cod)(Anjali et al., 2019; Cline, 2012; Muñoz-Colmenero et al., 2017; Xiong et al., 2016) or conservation concerns (Logan, Alter, Haupt, Tomalty, & Palumbi, 2008; Marín et al., 2018; Wainwright et al., 2018). Such standardised sampling would allow for a direct comparison across studies and regions. They would also allow for studying seafood mislabelling rates over time.

We would argue that the main problem with Singapore’s seafood products is ‘creative labelling’, especially for heavily processed products. This is likely due to the lack of clear regulations defining which species should be included in products carrying a particular common names. The Sale of Food Act (Cap. 283, RG 1) only states that labels need to provide a name or description which is “sufficient to indicate the true nature of the food”, as well as defining ‘fish’ as any aquatic organism commonly consumed by humans, excluding mammals, but explicitly including crustaceans and molluscs. This rules out egregious cases of mislabelling such as the use of pork in seafood products, but it allows for creative labelling. Arguably, this state of affairs is no longer in line with the expectation of today’s consumers who expect labels to be precise. This suggests that there may be a need for a regulatory update that could follow the example set by the European Union. The EU mandates that both the commercial and scientific name should be listed and that the commercial name be taken from approved lists published by EU member countries (Regulation (EU) No 1379/2013). The implementation of these rules resulted in a drop in the incidence of mislabelling of commercially sold seafood in EU supermarkets (from ca. 20% to ca. 8%: (Mariani et al., 2015), while countries with less strict laws continue to have mislabelling rates of about 20-30% (Carvalho et al., 2015; Hu et al., 2018; Nagalakshmi et al., 2016). Levels of seafood mislabelling may also drop in Singapore’s supermarkets if such legislation were to be enacted. Note, however, that the seafood mislabelling rates in Europe’s restaurants did not benefit from the new regulations (Christiansen, Fournier, Hellemans, & Volckaert, 2018; Horreo, Fitze, Jiménez-Valverde, Noriega, & Pelaez, 2019; Pardo et al., 2018), but this may not be a major concern in Singapore where all seafood samples obtained from restaurants were correctly labelled (N=21).

## 4. Conclusions

Our results suggest that MinION is ready for DNA-based monitoring for seafood. MinION reads can be used to identify key ingredients in single- and multi-species products even if they were heavily processed. We surmise that methods based on MS are likely to be the best choice for the routine identification of single-species samples of common species, but we would argue that DNA sequencing is the most suitable tool for mixed-species samples or samples of rare species lacking MS profiles. Developing better techniques for mixed-sample products is particularly important because some contain ingredients that should be highlighted on the labels while others appear to lack ingredients that are listed. Testing such samples can now be accomplished rapidly with MinION at a reasonable cost. The barcodes in our study still cost ca. USD 10 per sample, but this was an artefact of only sequencing 105 samples on one flowcell. The correct capacity is closer to 1000 samples (Srivathsan et al., 2019) even if two sets of primers are used. Fortunately, sequencing at smaller scales can also be cost-effective because flowcells can be used multiple times. Each re-use lowers the capacity which allows for having flowcells that are suitable for experiments of different sizes. In addition, small-scale projects can be carried out on new, lower-capacity flowcells (“Flongle”). Overall, we would thus predict that the consumable cost of MinION barcodes will be <USD1 per sample. Of course, implementing a fully developed monitoring scheme would require more than just a good sequencing method. It will require well-designed sampling methods, the development of explicit labelling guidelines, user-friendly bioinformatics software, and experimentally determined detection levels for ingredients in mixed-species samples.

## Acknowledgements

The authors would like to thank Yuen Huei Khee for help provided in the laboratory, as well as Allie Wharf and Freya Slessor (Make Waves Media) for their help obtaining samples. This work was supported by the South East Asian Biodiversity Genomics (SEABIG) Centre (Grant no: R-154-000-648-646 and R-154-000-648-733).

